# Inactivation of EMILIN-1 by proteolysis and secretion in small extracellular vesicles favors melanoma progression and metastasis

**DOI:** 10.1101/2021.06.02.446715

**Authors:** Ana Amor Lopez, Marina S. Mazariegos, Alessandra Capuano, Pilar Ximénez-Embún, Marta Hergueta-Redondo, Juan Ángel Recio, Eva Muñoz, Fátima Al-Shahrour, Javier Muñoz, Diego Megías, Roberto Doliana, Paola Spessotto, Héctor Peinado

## Abstract

Several studies have demonstrated that melanoma-derived extracellular vesicles (EVs) are involved in lymph node metastasis; however, the molecular mechanisms involved are not defined completely. Here, we found that EMILIN-1 is proteolyzed and secreted in small EVs (sEVs) as a novel mechanism to reduce its intracellular levels favoring metastasis in lymph node metastatic cells. Interestingly, we observed that EMILIN-1 has intrinsic *tumor and metastasis suppressive-like* properties reducing effective migration, cell viability, primary tumor growth and metastasis in mouse melanoma models. Finally, analysis in human melanoma samples showed that tumor cells with high levels of EMILIN-1 are reduced in metastatic lesions compared to primary tumors or nevi. Overall, our analysis suggests that the inactivation of EMILIN-1 by proteolysis and secretion in sEVs reduce its intrinsic *tumor suppressive* activities in melanoma favoring tumor progression and metastasis.

## Introduction

The tumor microenvironment has been found to play an active role in tumor progression [1]. Tumors induce the formation of microenvironments at distant organs that are conducive to the survival and outgrowth of tumor cells prior to their arrival at these sites. These microenvironments were termed “pre-metastatic niches” (PMNs) [2]. This concept proposes the ability of primary tumor cells to precondition regional and distal organs for future metastatic disease before the arrival of circulating tumor cells via tumor-derived factors. Therefore, PMN represents an abnormal and favourable microenvironment for metastasis [2].

Several reports have highlighted the role of extracellular vesicles (EVs) during PMN formation [3, 4]. EVs are composed by a lipid bilayer that contains molecular cargo representative from the cell of origin (e.g., proteins, RNA, DNA, etc.) [5]. EVs mediate cell-cell communication by several mechanisms, from inducing intracellular signaling after their interaction, to confer new properties due to the acquisition of new receptors, enzymes or even genetic material after their uptake [6]. A recent classification based on size divided them in large EVs (lEVs) and small EVs (sEVs) [7]. Microvesicles (200 nm - 1μm), apoptotic bodies (1-5 μm) and oncosomes (1-10 μm) are considered lEVs [7, 8]. sEVs serve as a vehicle for horizontal transfer of molecules such as RNAs, DNA and proteins which, once in the target cell, can exert their function [4–6, 9, 10].

Tumors induce changes in the sentinel LN (LNs) such as enhanced lymphangiogenesis [11, 12], induction of an immunosuppressive environment and increased vascular flow [13, 14]. These changes precede metastatic colonization and contribute to the formation of the PMN in the LNs [11, 13, 14]. Understanding the mechanisms involved in LN metastasis is crucial to define the first steps of melanoma metastatic spread. Given that sEVs-derived from melanoma cells promote lymphangiogenesis, extracellular matrix (ECM) remodeling, immunosuppression and metastasis in LNs [15, 16], in this work we wanted to analyze if lymph the node metastatic melanoma model B16-F1R2 has a specific signature that may favor tumor cell survival in lymph nodes. We have found a specific signature of genes over-expressed in cells and proteins hyper-secreted in sEVs including EMILIN-1, a protein involved in lymph node physiology and pathology [17, 18]. We found that EMILIN-1 is proteolyzed and secreted in sEVs as a mechanism to reduce its intracellular levels in this cell line. In order to analyze the relevance or our findings, we overexpressed EMILIN-1 in B16-F1 cells and found a reduction of effective migration and cell viability suggesting that EMILIN1 has intrinsic *tumor suppressive* activities in melanoma. Noteworthy, EMILIN-1 overexpression led to reduced primary tumor and lymph node metastasis in B16-F1 mouse melanoma xenograft models. Finally, we found that cells expressing high levels of EMILIN-1 are reduced in human melanoma metastatic lesions compared to primary tumors and nevi. Overall, our data suggest that the proteolysis of EMILIN-1 and its secretion in sEVs is a novel mechanism of EMILIN-1 inactivation favoring melanoma metastasis.

## Results

### Characterization of secreted sEVs in melanoma models

We have characterized secreted sEVs from a panel of mouse melanoma models representative of low metastatic potential (B16-F1), high metastatic potential (B16-F10), and lymph node metastasis (B16-F1R2) [19] **(Fig.1A).** We first isolated by ultracentrifugation and characterized secreted sEVs from the melanoma models described. Analysis of protein in secreted sEVs demonstrated that, congruently with our previous published data [26], the lymph node metastatic cell line (B16-F1R2) and the high metastatic model (B16-F10) secrete increased protein cargo in sEVs **(Fig.1B, C)** than the poorly metastatic cell line B16-F1. Analysis by nanoparticle tracking analysis showed a typical distribution size of sEVs (**Supplementary Figure 1**).

**Fig.1.**
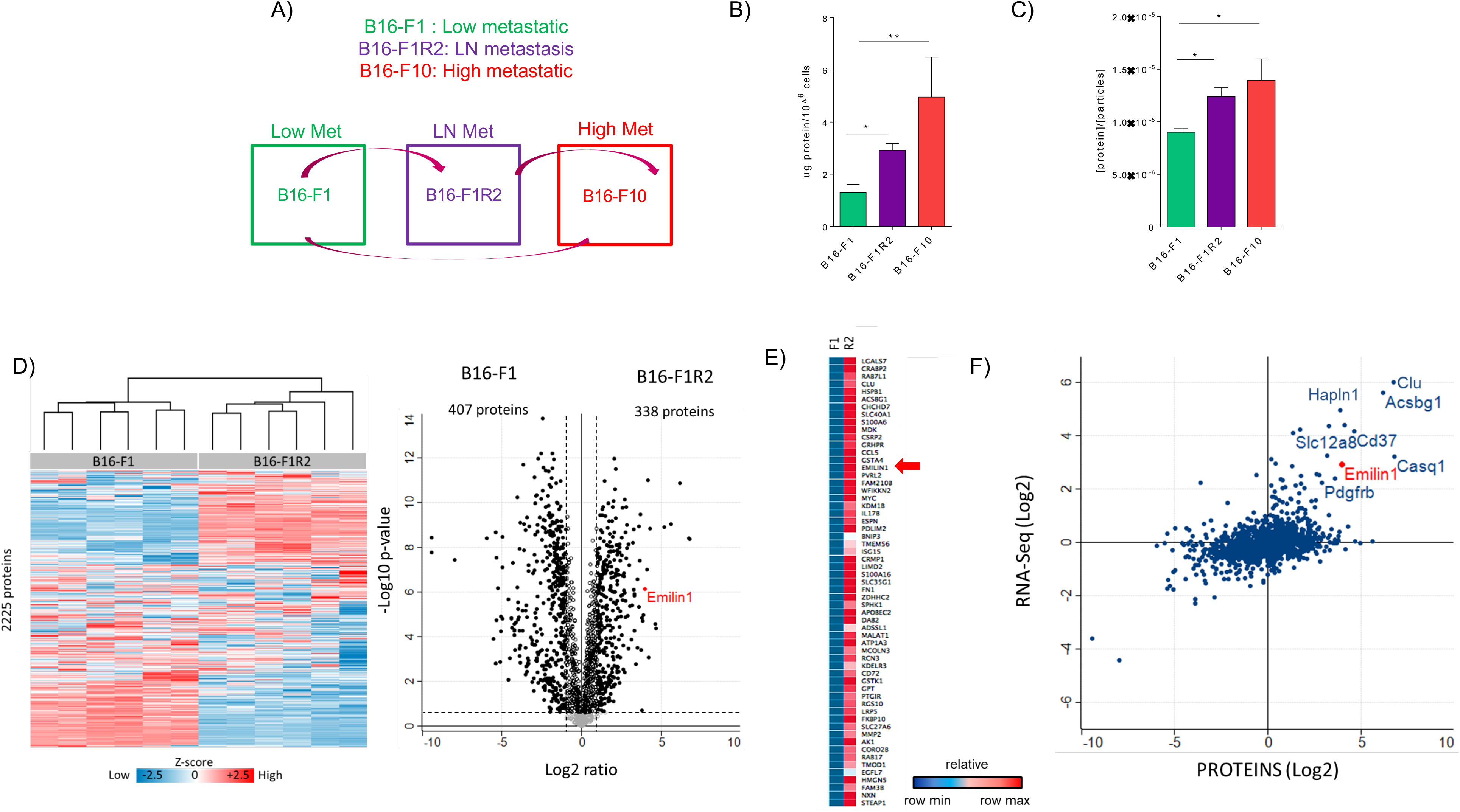
Defining the proteomic signature of sEVs secreted from lymph node metastatic melanoma models. A) Schematic representation of the mouse melanoma models used. B) Analysis of sEV number secretion and C) total protein secreted in sEVs in mouse melanoma cells lines derived from low metastatic model (green), lymph node metastasis (purple) and distal metastasis (dark purple and red). n=5 . p <0.05 using Mann Whitney test. D) Unsupervised hierarchical clustering. 2225 proteins were identified in sEVs derived from B16-F1 and B16-F1R2 cell lines. To the right, a volcano plot showing the differentially expressed proteins (Student’s t test, p < 0.05). Black circles represent proteins above the fold change cutoff. E) Heatmap of genes differentially expressed in B16-F1R2 and –R2L compared to B16-F1. As noted by the arrow, EMILIN-1 is over-expressed in both cell lines. F) Correlation of proteomic and transcriptomic data to define the main genes overexpressed and proteins hyper-secreted in sEVs from B16-F1R2 compared to B16-F1 cell line.

To investigate the proteomic signatures in the sEVs associated to the lymph node metastatic melanoma model B16-F1R2, we collected sEVs from this model and the parental B16-F1 model and performed mass spectrometry analysis **(Fig.1D).** We identified 2225 proteins in the sEVs derived from these models and confirmed that samples clustered by cell line after performing unsupervised hierarchical clustering analysis **(Fig.1D).** Differential analysis showed that 33% of the proteins were significantly regulated. Among them, we found 338 upregulated proteins in B16-F1R2–derived sEVs compared to parental-derived sEVs suggesting the existence of a proteomic signature associated to the LN metastatic model. Specifically, we found an enrichment in proteins related ECM to organization **(Supplementary Fig. 2).**

We next correlated the protein cargo of secreted sEVs with gene expression analysis in B16-F1 and B16-F1R2. We performed RNA sequencing analysis of B16-F1, B16-F1R2, we selected a subset of the top genes most upregulated in B16-F1R2 model compared to B16-F1 **(Supplementary Table 1, example of cluster Fig. 1E).**

We observed that some overexpressed genes belonged to proteins hyper-secreted in sEVs from B16-F1R2; therefore, we integrated the mass spectrometry data (proteins secreted in sEVs) with the RNA sequencing data. Interestingly, out of several candidates we selected EMILIN-1, a protein related to ECM and lymph node remodeling, that was hyper-secreted in sEVs and also overexpressed at the mRNA level (**Fig.1F, Supplementary Table 2**).

### EMILIN-1 is proteolyzed and secreted in sEVs from B16-F1R2 cell line

Elastin microfibrillar interface proteins (EMILINs) constitute a four-member family of glycoproteins with a C-terminus gC1q domain typical of the gC1q/TNF superfamily members and also contain N-terminus unique cysteine-rich EMI domain [27]. EMILIN-1 has been characterized in multiple scenarios such as cell migration [28] and proliferation [27], lymphatic vessel function [19] skin homeostasis [27] and cancer development [26]. Analysis of EMILIN-1 expression by qPCR showed that while it is highly expressed in melanocytes (melan-a), its levels were downregulated level in B16-F1 and then re-expressed in B16-F1R2 and F10 at mRNA **(Fig.2A).** Importantly, analysis of EMILIN-1 expression by Western-blot using specific antibodies [26] in mouse melanoma models demonstrated that it is not detected intracellularly in none of the tested cell lines but interestingly it is secreted in sEVs derived from LN metastatic model B16-F1R2 **(Fig.2B, left panels, cells),** which let us to hypothesize that EMILIN-1 could be detected extracellularly. Indeed, supporting our mass spectrometry data, EMILIN-1 was detected in sEVs but with several bands with a lower molecular weight than expected (150 KDa) as opposed to the full-length protein used as a positive control **(Fig.2B, right panels, sEVs)**, suggesting its proteolysis. Interestingly EMILIN-1 has been previously reported to be proteolyzed in other models [29, 30], suggesting that secretion and proteolysis of this protein in melanoma cells could be a novel mechanisms of inactivation.

**Fig.2.**
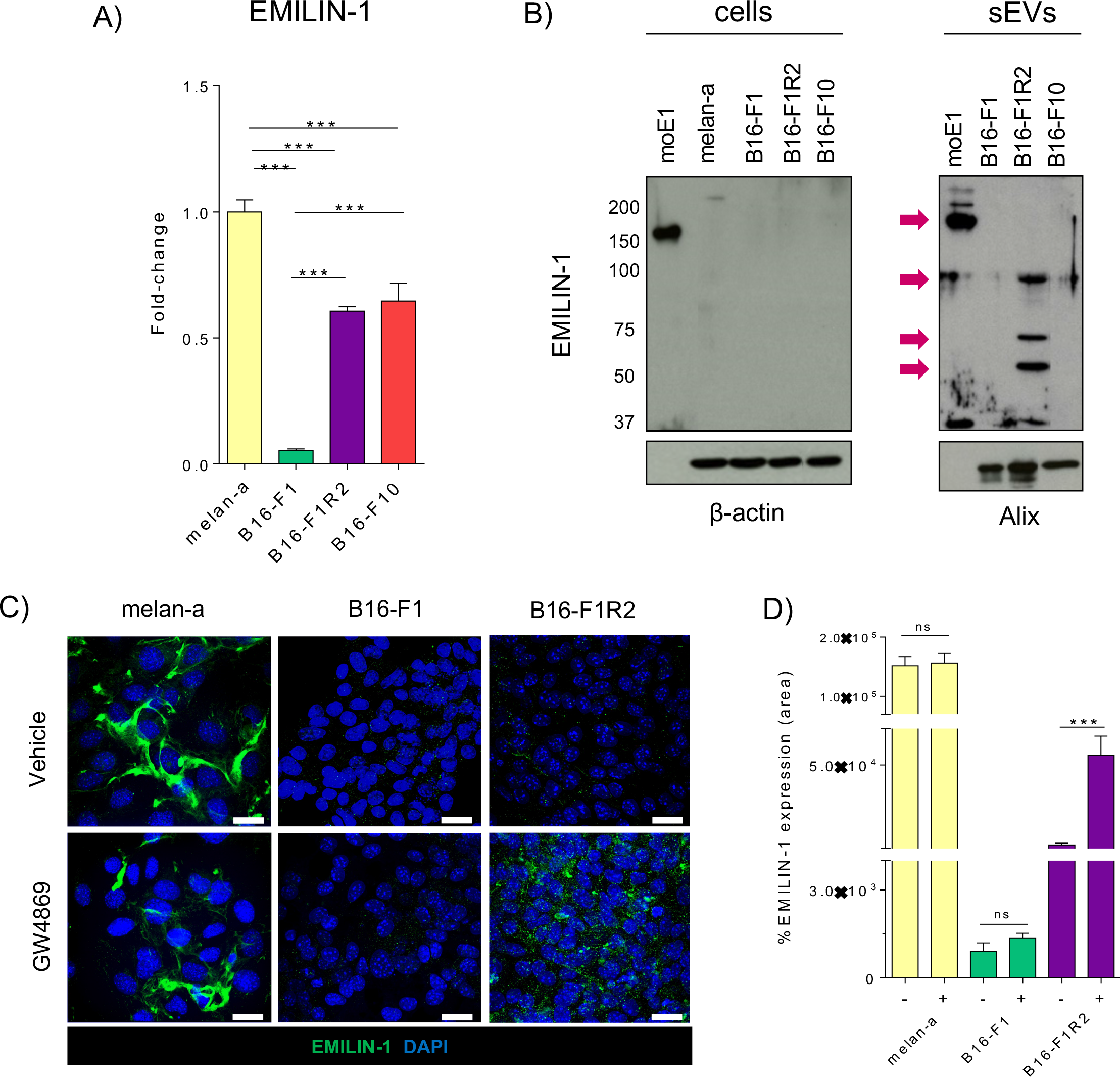
EMILIN-1 is degraded and secreted in sEVs from B16-F1R2 cell line. A) mRNA expression levels by qRT-PCR of EMILIN-1 (n=2). B) Analysis by Western-blot of EMILIN-1 in mouse cell line models and derived sEVs (B16-F1, B16-F1R2, B16-F1R2L and B16-F10). moE1, EMILIN-1 recombinant mouse protein, was used as loading positive control, molecular weight expected is 150 kDa indicated by an arrow. Anti-EMILIN-1 antibody detected several bands with a lower molecular weight than expected in secreted sEVs, indicated by an arrow. C) Analysis of EMILIN-1 (in green) expression and localization by confocal immunofluorescence (scale 18 μm) in melanocytes, B16-F1 and B16-F1R2 cell lines before and after the treatment with 10 μM GW4869 during 24 hours. Cell nuclei were stained with DAPI (in blue). D) Quantification of EMILIN-1 expression (green signal) in C), *p<0,05 by Non-parametric t-test.

We next decided to analyze if EMILIN-1 secretion in sEVs could be inhibited by the use of the EV secretion inhibitor GW4869, a non-competitive inhibitor of sphingomyelinase. In agreement with WB and RNA expression data, we observed that EMILIN-1 is expressed and deposited in the ECM by melan-a cell line, however is not detected in B16-F1 and B16-F1R2 cells **(Fig. 2C, upper panels).** Treatment with GW4869 did not affect EMILIN-1 in melan-a and B16-F1 cells **(Fig.2C, lower left and middle panels)**, suggesting that extracellular deposition of EMILIN-1 by melan-a cells is not mediated by EV secretion. However, we observed a significant accumulation of EMILIN-1 in B16F1-R2 cell line after treatment with GW4869 **(Fig.2C, lower right panel)**. We performed similar studies but using the proteasome inhibitor MG-132, a potent, reversible, and cell-permeable proteasome inhibitor and reduces the degradation of ubiquitin-conjugated proteins in mammalian cells [31, 32]. We observed that Emilin1 levels remained similar after MG-132 treatment in all cell types, suggesting that proteasome inhibition does not affect Emilin1 levels. **(Supplementary Fig. 3).** These data show that inhibition of EV secretion avoid EMILIN-1 secretion extracellularly in B16-F1R2 cell line.

### EMILIN-1 overexpression reduces cell viability and effective migration

Since EMILIN-1 has been already reported to have tumor suppressive like functions [18, 33], we postulated that melanoma cells secrete and degrade EMILIN-1 in sEVs as a novel mechanism to inactivate its suppressive signals in melanoma. To define the intrinsic role of EMILIN-1 in melanoma cells, we performed cell viability and cell cycle assays. We first analyzed cell viability in B16-F1-HA control cells or cells overexpressing HA-EMILIN-1 (B16-F1-HA-E1). To define the effects on cell proliferation we analyzed number of viable cells at different time points (24, 48 and 72h). We observed that overexpression of EMILIN-1 reduced significantly B16-F1 cell viability in three different clones analyzed **(Fig.3A).** To determine if this reduction was due to changes in cell cycle, we analyzed cell cycle histograms for bulk DNA staining (PI), after addition of EdU, in B16-F1-HA and B16-F1-HA-E1 cells after 24h, 48h, 72h, 96 h **(Fig.3B).** We found that EMILIN-1 overexpression does not affect significantly cell cycle in B16-F1 cells. These results suggest that Emilin1 affect mainly the metabolic balance and cell viability in cells but not the cell cycle of the cells. Analysis of the percentage of cells in S phase was then calculated. We found that EMILIN-1 overexpression does not affect significantly cell cycle compared to B16-F1-HA control cells **(Fig.3C).** Then, we performed melanoma cell migration tracking assays in B16-F1-HA and B16-F1-HA-E1 cells. We observed in their trajectory reconstruction that there is a loss of migration directionality in cells overexpressing EMILIN-1 compared to control cells B16-F1-HA cells that had a defined migration **(Fig.3D).** These data suggest that increased EMILIN-1 levels reduce effective migration in melanoma cells. Therefore, it is plausible that reduction of intracellular EMILIN-1 levels observed along melanoma progression could be required for an efficient and independent migration, a well-known property of melanoma cells.

**Fig.3.**
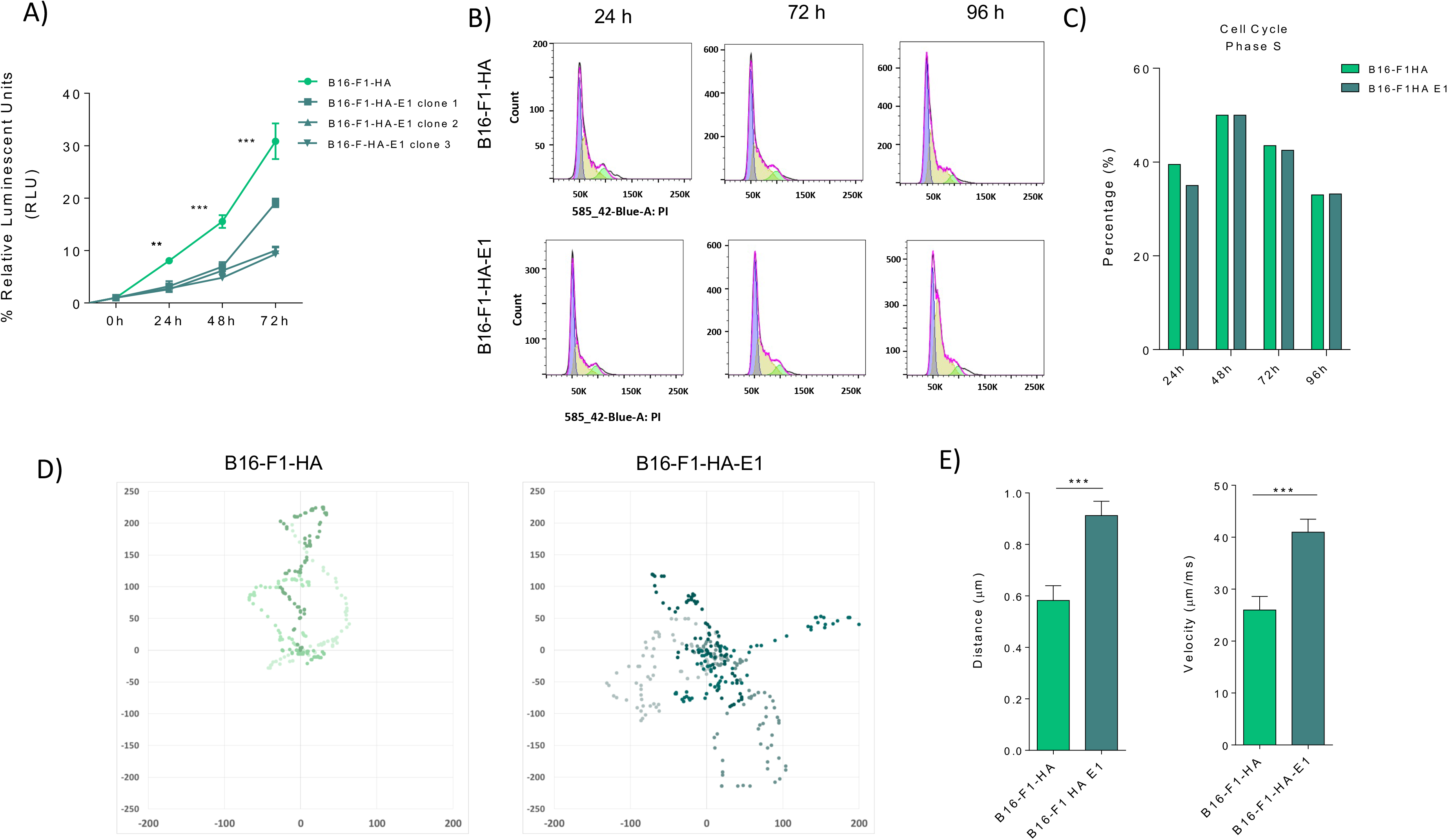
EMILIN-1 Emilin1 overexpression reduces cell viability and directionality cell migration. A) Analysis of cell viability using CellTiter-Glo® Luminescent Cell Viability Assay after EMILIN-1 expression of B16-F1-HA and different clones of B16-F1-HA cells overexpressing EMILIN-1 at indicated time point (n=2) **p<0.01 and ***p<0.001 by 2-way ANOVA. Relative Luminescent Units (RLU) are represented in y axis. B) Representative histograms showing the cell cycle distribution in B16-F1-HA and B16-F1-HA-E1 models at 24, 72 and 96 hours. C) Relative changes in the percentage of B16-F1-HA and B16-F1-HA-E1 cells in S phase after addition of Edu. D) Representative examples of cell tracking and motility analysis in B16-F1-HA and B16-F1-HA-E1 models. E) Distance and velocity of cell tracking analysis by ImageJ software ***p<0,001 by Unpaired t-test

### EMILIN-1 overexpression reduces primary tumor growth and lymph node metastasis

We next analyzed the effect of EMILIN-1 overexpression in tumor growth and metastatic properties of B16-F1 cells. For this purpose, C57BL/6 mice were injected subcutaneously with B16-F1-HA control cells or cells overexpressing EMILIN-1 (B16-F1-HA-E1). We found that EMILIN-1 overexpression led to a significant reduction of tumor growth **(Fig.4A).** Moreover, analysis of lymph node metastasis after intra-footpad injection showed a reduction of lymph node metastasis after EMILIN-1 overexpression **(Fig.4B, 4C).** These data support that EMILIN-1 overexpression reduces both primary tumor growth and lymph node metastasis in B16-F1 cells.

**Fig.4.**
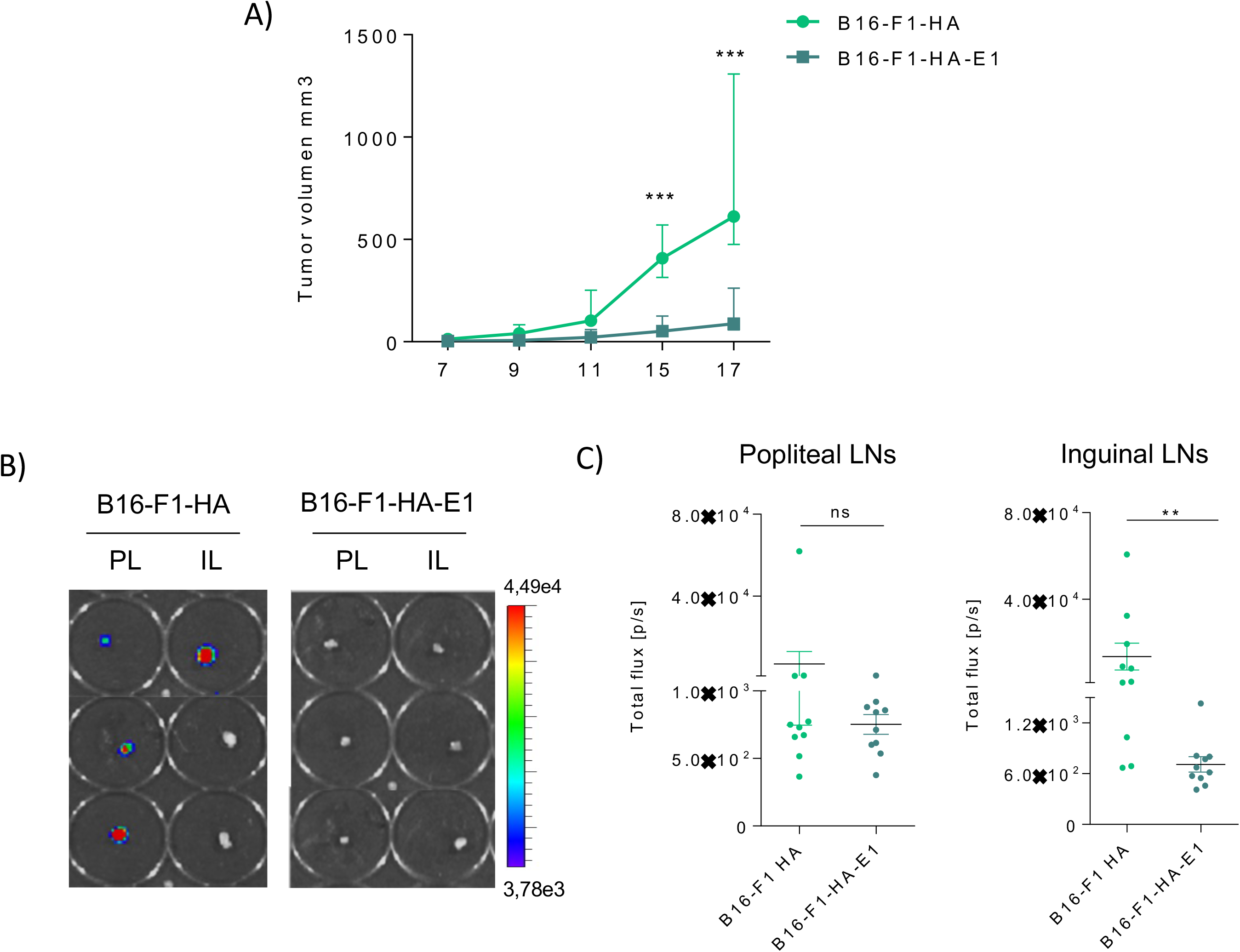
Overexpression of EMILIN-1 reduce tumor growth and lymph node metastasis. A) Growth of subcutaneous xenografts established from B16-F1 and B16-F1-HA E1 cells (n=5 per group) ***p<0.001 by 2-way ANOVA. B) In vivo imaging of popliteal and inguinal lateral LNs, from intrafootpad injected mice from B16-F1-HA andB16-F1-HA-E1 cells. C) Total flux (p/s, photons per second) quantification of popliteal and inguinal LN, n=10**p<0.05 by Nonparametric t-test

### EMILIN-1 stabilization leads to reduced lymph node metastasis

Among all the proteolytic enzymes released by the tumor, neutrophil elastase (NE) was found as the main enzyme able to fully impair the regulatory function of EMILIN1 in sarcoma and ovarian cancer [18, 33]. The consequence of this proteolytic process was the impairment of its anti-proliferative role [29, 30]. The local administration of sivelestat, an inhibitor of neutrophil elastase prevents EMILIN1 degradation and reduces lymphoedema, restoring a normal lymphatic functionality in a mouse lymphoedema model [29, 30]. Analysis by site direct mutagenesis in 914 residue of EMILIN1 demonstrated that mutant R914W was resistant to NE proteolytic cleavage [29, 30]. Based on this, we next analyzed the effect of the EMILIN-1 stabilization mutant (E1-R914W) [29, 30] in B16-F1R2 cells in primary tumor growth and metastasis. We observed that while EMILIN-1 stabilization mutant did not affect significantly primary tumor growth **(Fig.5A, 5B)**, it reduced significantly spontaneous lymph node metastasis **(Fig.5C, 5D).** We also analyzed the effect in lymph node experimental metastasis after footpad injection and we observed that EMILIN-1 stabilization mutant expression reduced lymph node metastases **(Fig. 5E, 5F, 5G)**.

**Fig.5.**
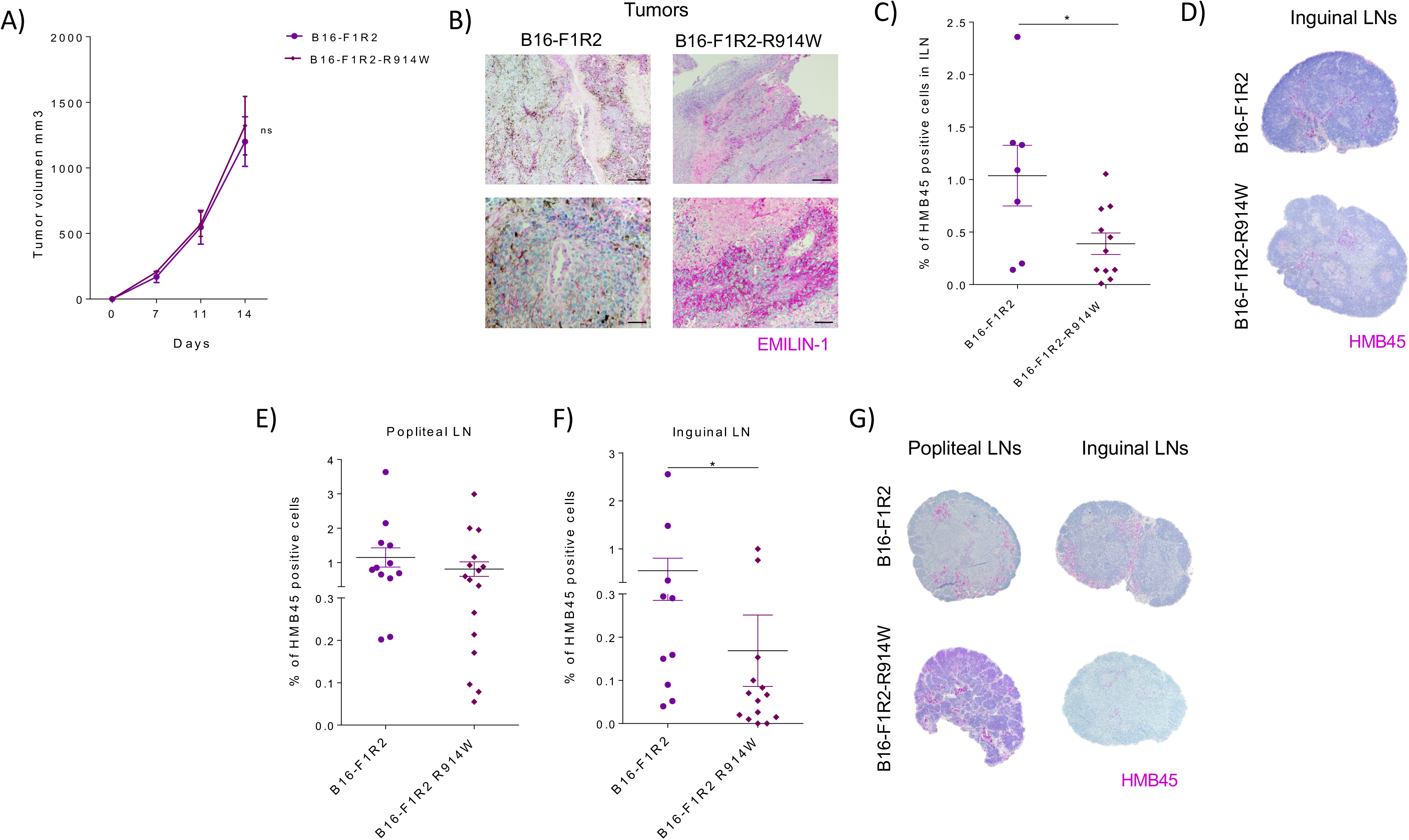
Analysis of EMILIN-1 stabilization mutant in vivo. A) Tumor growth and lymph node metastasis of B16-F1R2 and B16-F1R2-R914W xenografts. C57BL/6 mice were injected subcutaneously with 1×10^6^ B16-F1R2 (n=8 mice) and B16-F1R2-R914W (n=10 mice) cells. B) Representative images of B16-F1R2 and B16-F1R2-R914W tumor sections stained for EMILIN-1. Bar: 100μm C) Percentage of melanoma HMB45 positive cells quantification from inguinal lymph nodes were used for analyzed lymph node metastasis (B16-F1R2, n=7 and B16-F1R2-R914W, n=13) *p<0.05 by Nonparametric ttest. D) Representative images of inguinal lymph nodes (HMB45 staining) from B16-F1R2 and B16-F1R2 R914W xenografts. Bar: 5001m. E-G) Analysis of lymph nodesfrom C57BL/6 mice injected intrafootpad with 200,000 B16-F1R2 and B16-F1R2-R914W cells. Percentage of Melanoma HMB45 positive cells quantification from E) popliteal (B16-F1R2 group n=12 and B16-F1R2-R914W group n=16) and F) inguinal lymph nodes (B16-F1R2 group n=10 and B16-F1R2-R914W group n=14), *p<0,05 by Nonparametric t-test. G) Representative images of popliteal and inguinal lymph nodes (HMB45 staining) from C57BL/6 mice. Bar: 500μm

## Discussion

During the last years, several studies have demonstrated that EVs play an important role in cell-cell communication being actively involved in tumor progression and metastasis [34]. Data support that tumor-derived EVs are key players in the formation of pre-metastatic niche formation at distal sites and metastatic organotropism [2]. Consistent with this, tumor-derived sEV can reach the sentinel lymph nodes favoring metastatic spread [15, 16]. Indeed, they have been proposed to play a role in lymph node pre-metastatic niche formation [11]. It has been previously reported that metastatic melanoma models secrete higher amounts of proteins in sEV as compared to low metastatic models [26]. In addition, sEV protein concentrations are higher in the plasma and seroma of melanoma subjects with higher stages (III, IV) compared to lower stages (I, II) and subjects without cancer [26, 35]. In agreement with that, we observed that sEVs from lymph node metastatic (B16-F1R2) and high metastatic models (B16-F10) secrete higher amounts of protein in sEV than poorly metastatic cell line B16-F1. However, the gene expression signatures associated to melanoma lymph node metastatic cells on sEV have never been defined. Therefore, in this work, we have characterized the gene expression profile and the proteomic signature associated to sEV in the lymph node metastatic model B16-F1R2 [19]. Analysis of the mRNAs over-expressed in the cells and proteins hyper-secreted in sEV showed a signature related to ECM and processes linked to tumor-microenvironment interaction. Among all the proteins observed in the signature, we focused our analysis on EMILIN-1 due to its relevance in processes such as ECM remodeling, cell adhesion, lymphatic vessel functionality and proliferation [18, 30, 36–38].

Importantly, we found that EMILIN-1 was not detected at protein level intracellularly in B16-F1R2, but seems to be proteolyzed and secreted through sEV, suggesting that sEV contribute to the clearance of the protein in tumor cells. Our data show that EMILIN-1 is proteolyzed and secreted in sEV while is still detected at mRNA level but undetectable in sEV from the highly metastatic model B16-F10. These results suggest that EMILIN-1 proteolysis and secretion in sEV favor lymph node metastasis, while additional mechanisms of EMILIN-1 inactivation (e.g., reduction of protein half-life or translational repression) may arise later in highly metastatic models. Interestingly, some studies have suggested that expulsion of specific cargo in sEVs may be a mechanism of eliminating tumor suppressor molecules (e.g., miRNAs, proteins) favoring survival of cancer cells [39–41]. Therefore, we postulated that EMILIN-1 secretion and degradation in sEVs might be a novel mechanism involved in EMILIN-1 inactivation melanoma progression and metastasis.

Indeed, expression of EMILIN-1 has already been described with suppressive activities in epithelial cells, colitis and colon carcinogenesis [17, 42]. Importantly, analysis in EMILIN-1 KO showed that its ablation in the microenvironment promoted tumor progression in skin, melanoma and colon cancer models [18]. EMILIN-1 degradation is already reported, for example by neutrophil elastase (NE), a specific mechanism leading to the loss of functions disabling its tumor suppressor-like properties [33]. Using a proteomic approach, EMILIN-1 has been defined as a potential candidate substrate for MMP-3, −9 and MT1-MMP in the cardiovascular system [43]. In this study, we found that EMILIN-1 secreted in sEVs is proteolyzed in a similar pattern than previously described [29]. However, we used NE and MMP inhibitors in vitro in our models but we did not observe any effect (data not shown), suggesting that these protease activities are not involved in EMILIN-1 proteolysis in B16-F1R2 cells. Defining the EMILIN-1 “degradome” in specific melanoma scenarios could be crucial to find novel inhibitors that may lead to the reactivation of its suppressive activity in vivo. Interestingly, we analyzed the effect of the extracellular vesicle secretion inhibitor GW4869 on the regulation of EMILIN-1 and found that it is accumulated intracellularly after inhibition demonstrating its active secretion in sEVs .

The fact that EMILIN-1 is lost along melanoma progression, suggest its role as a tumor suppressor-like protein. Hence, we studied the intrinsic role of EMILIN-1 in melanoma cells by analyzing its influence in cell viability and cell cycle assays. We observed that EMILIN-1 overexpression in B16-F1 cell line reduced cell viability. These results are in agreement with published data defining that EMILIN-1 expression in the ECM reduces cell proliferation in normal and tumor cells [18, 29, 33]. Our analysis of cell cycle could not define any effect on it, but rather decrease in cell viability suggesting a metabolic activity. We also analyzed the impact of EMILIN-1 in cell migration. We observed that cells overexpressing EMILIN-1 had no directed migration, suggesting that its overexpression impair effective migration in melanoma cells. Previous analysis of Emillin1 expression in the skin showed that it locates in the dermis, up to the basement membrane, interacting with components of the ECM but also with the anchoring complex suggesting an important role for cell adhesion, migration, proliferation [45]. Directional cell motility is critical during inflammation, embryogenesis, and cancer metastasis [46, 47].

Notably, we observed a significant decrease in tumor growth in B16-F1 cells overexpressing EMILIN-1. Importantly, analysis of experimental lymph node metastasis after footpad injection showed a significant decrease in both models B16-F1R2-R914W and B16-F1-HA-E1. These data support that besides a tumor suppressor like protein, EMILIN-1 has an important effect reducing lymph node metastasis. Analysis of the main genes affected after EMILIN-1 overexpression may be interesting to define the mechanism involved in metastasis suppression.

Altogether, our data support that EMILIN-1 proteolysis and secretion in sEVs reduce its *tumor- and metastatic suppressive-like* properties favoring cell viability, effective migration, tumor growth and lymph node metastasis in mouse melanoma cells. Overall, our data support a novel and not previously defined intrinsic tumor suppressive activity of EMILIN-1 in melanoma cells that is abolished by its proteolysis and secretion in sEVs favoring tumor progression and metastatic behavior.

## Acknowledgements

We thank the members of Dr Peinado’s laboratory for helpful discussions. The authors gratefully acknowledge the support of the following sources of funding: MINECO (SAF2014-54541-R), Ramón y Cajal Programme, Asociación Española Contra el Cáncer, La Caixa Foundation and FERO Foundation. We are also grateful for the support of the MINECO-Red de Excelencia TENTACLES and MINECO-Severo Ochoa predoctoral program.

## Authors contributions

A. A. L., M. S. M., A. C. M. H.-R, performed experiment and data analysis P.X-E, J. M. F. Al-S performed data analysis, J. Á. R. and E. M. were involved in analysis of human samples, D. M. helped with miccroscopy analysis, R. D., P. S. contributed to the scientific discussion of the work and H. P. conceived and directed the study. A. A. L and H. Planed the experiments and wrote the manuscript. All authors contributed to and approved the final version of the manuscript.

## Ethics declarations

The authors declare no competing financial interests.

## Methods

### Cell lines

B16-F1 and B16-F10 were purchased from American Type Culture Collection (ATCC). The lymph node metastatic variant B16-F1R2 and B16-F1R2L was kindly provided by Dr. Michael Detmar and Dr. Steven Proulx (ETH Zurich, Switzerland) [19]. Spontaneously immortalized mouse melanocytes cell line (melan-a) were kindly provided by Dr. Dorothy C Bennett, (St. George’s University of London). All melanoma cell lines were grown in high glucose DMEM (Lonza #D6429) supplemented with 10% fetal bovine serum (Hyclone #SH30071.03IH), 2 mM glutamine and 20 μg/ml gentamicin (Sigma #G1272). melan-a cell line was cultured in RPMI (Gibco #11875-093), supplemented with 5% fetal calf serum (Gibco #16030074) and 200 nM 12-Otetradecanoyl phorbol-13-acetate (Sigma #16561-29-8). All cells were grown at 37°_C in a humidified 5% CO_2_ atmosphere and routinely tested for mycoplasma contamination.

### sEVs purification

Cells were cultured in medium supplemented with 10% sEV-reduced fetal bovine serum (Hyclone#SH30071.03IH). FBS was reduced from bovine sEVs by ultracentrifugation at 100,000xg for 70 min. For sEV isolation, conditioned medium for 72 hours was centrifuged at 500xg for 10 min RT to remove cell contaminants. Then, to remove big debris and microvesicles the supernatant fraction was centrifuged at 12,000xg for 20 min at 10°_C. SEVs were then harvested by ultracentrifugation at 100.000xg for 70 min. The sEVs pellet was washed in 20 ml of PBS and sEVs were collected by ultracentrifugation at 100,000xg for 70 min. All ultracentrifuge spins were performed at 10°_C using a BECKMAN Optima X100 centrifuge with BECKMAN TYPE 70.1Ti rotor. Final sEVs pellet was resuspended in 100 μL of PBS and the protein content was measured by bicinchoninic acid assay (Pierce™ BCA Protein Assay Kit, Thermo Scientific). The NS500 nanoparticle characterization system (NanoSight) equipped with a blue laser (405 nm) was used for real-time characterization of the vesicles.

### Sample preparation for proteomic analysis

Proteins were solubilized using 8 M urea in 100 mM Tris-HCl pH 8.0. Samples (10 μg) were digested by means of the standard FASP protocol (43). Briefly, proteins were reduced and alkylated (15 mM TCEP, 30 mM CAA, 30 min in the dark, RT); and sequentially digested with Lys-C (Wako) (protein:enzyme ratio 1:50, o/n at RT) and trypsin (Promega) (protein:enzyme ratio 1:50, 6 h at 37 °C). Resulting peptides were desalted using C18 stage-tips.

### Mass spectrometry

LC-MS/MS was done by coupling a nanoLC-Ultra 1D+ system (Eksigent) to a LTQ Orbitrap Velos mass spectrometer (Thermo Fisher Scientific) via a Nanospray Flex source (Thermo FisherScientific). Peptides were loaded into a trap column (NS-MP-10 BioSphere C18 5 μm, 20 mm length, NanoSeparations) for 10 min at a flow rate of 2.5 μl/min in 0.1% FA. Then peptides were transferred to an analytical column (ReproSil Pur C18-AQ 2.4 μm, 500 mm length and 0.075 mm ID) and separated using a 120 min linear gradient (buffer A: 4% ACN, 0.1% FA; buffer B: 100% ACN, 0.1% FA) at a flow rate of 250 nL/min. The gradient used was: 0-2 min 6% B, 2-103 min 30% B, 103-113 min 98% B, 113-120 min 2% B. The peptides were electrosprayed (1.8 kV) into the mass spectrometer with a PicoTip emitter (360/20 Tube OD/ID μm, tip ID 10 μm) (New Objective), a heated capillary temperature of 325°_C and S-Lens RF level of 60%. The mass spectrometer was operated in a data-dependent mode, with an automatic switch between MS and MS/MS scans using a top 15 method (threshold signal ≥ 800 counts and dynamic exclusion of 45 sec). MS spectra (350-1500 m/z) were acquired in the Orbitrap with a resolution of 60,000 FWHM (400 m/z). Peptides were isolated using a 1.5 Th window and fragmented using collision induced dissociation (CID) with linear ion trap read out at a NCE of 35% (0.25 Q-value and 10 ms activation time). The ion target values were 1E6 for MS (500 ms max injection time) and 5000 for MS/MS (100 ms max injection time). Samples were analyzed twice.

### Proteomic data analysis

Raw files were processed with MaxQuant (v 1.5.3.30) using the standard settings against a human (UniProtKB/Swiss-Prot, August 2014, 20,187 sequences) or mouse (UniProtKB/Swiss-Prot/TrEMBL, August 2014, 43,539 sequences) protein database, supplemented with contaminants. Label-free quantification was performed with match between runs (match window of 0.7 min and alignment window of 20 min). Carbamidomethylation of cysteines was set as a fixed modification whereas methionine oxidation and N-term acetylation were variable protein modifications. The minimal peptide length was set to 7 amino acids and a maximum of two tryptic missed-cleavages were allowed. The results were filtered at 0.01 FDR (peptide and protein level). Afterwards, the “proteinGroup.txt” file was loaded in Perseus (v1.5.1.6) for further statistical analysis. A minimum of four LFQ valid values per group was required for quantification. Missing values were imputed from the observed normal distribution of intensities. Then, a two-sample Student’s T-Test with a permutation-based FDR was performed. Only proteins with a q-value<0.1 and log2 ratio >1 or < −1 were considered as regulated. The mass spectrometry proteomics data have been deposited to the ProteomeXchange Consortium via the PRIDE partner repository with the dataset identifier PXD018891. The following username and password (nHbVMAdB) can be used during the peer review purposes.

### RNA sequencing (RNA-Seq) and bioinformatics analysis

Total RNA was isolated from cells using the RNeasy Mini Kit (Qiagen #74106). The quantity and quality of the extracted RNA was assessed using NanoDrop ND-1000 Spectrophotometer (Thermo Scientific) and Agilent 2100 Bioanalyzer. RNA sequencing (RNA-seq) was performed by the CNIO Genomics Unit. 1 μg of total RNA from each sample was used. PolyA+ fraction was purified and randomly fragmented, converted to double stranded cDNA and processed through subsequent enzymatic treatments of end-repair, dA tailing, and ligation to adapters as in Illumina’s “TruSeq Stranded mRNA Sample Preparation Part # 15031047 Rev. D” kit (this kit incorporates dUTP during 2nd strand cDNA synthesis, which implies that only the cDNA strand generated during 1st strand synthesis is eventually sequenced). Adapter-ligated library was completed by PCR with Illumina PE primers (8 cycles). The resulting purified cDNA library was applied to an Illumina flow cell for cluster generation and sequenced using the Illumina HiSeq2500 platform by following manufacturer’s protocols. 50bp single-end sequenced reads were analyzed with the nextpresso pipeline [20] as follows: sequencing quality was checked with FastQC v0.10.1 (http://www.bioinformatics.babraham.ac.uk/projects/fastqc/). Reads were aligned to the mouse genome (NCBI37/mm9) with TopHat-2.0.10 [21] using Bowtie 1.0.0 [22] and Samtools 0.1.1.9 [23] allowing two mismatches and 20 multihits. Differential expression was calculated with DESeq2 [24], using the human NCBI37/mm9 transcript annotations from https://ccb.jhu.edu/software/tophat/igenomes.shtml. GSEAPreranked [25] was used to perform gene set enrichment analysis of the described gene signatures on a pre-ranked gene list, setting 1000 gene set permutations. Only those gene sets with significant enrichment levels (FDR q-value < 0.1) were finally considered. Access to RNA-seq data is provided from the Gene Expression Omnibus, under the ID GSE150221.

### Proteomic and RNAseq integration

We integrated the profiles associated to lymph node metastatic mouse model (B16-F1 vs B16-F1R2) data from RNAseq and proteomic analysis. Results were represented as the correlation of the ratios at the protein and at the RNAseq level (in log2).

### Gene expression analysis / Quantitative real-time PCR analysis

Cell lines were analyzed for specific gene expression using pre-designed primers listed below: EMILIN-1 (Fw 5’-CCTGTCTGGCTCCAGTGC-3’, Rv 5’-GCTCTAGCTGCTGCACCTTC-3’) and Hprt (Fw 5’-TCCTCCTCAGACCGCTTTT-3’, Rv 5’-CCTGGTTCATCATCGCTAATC-3’). In brief, total RNA was extracted from tissues or cells as described above and reverse-transcribed using the QuantiTect Reverse Transcription Kit (Qiagen #205313). Quantitative real-time PCR (QRT-PCR) was performed on a 7500 Fast Real Time PCR System (Applied Biosystems), using Sybergreen Universal PCR Master Mix (Life Technologies #4304437). Gene expression was analyzed using the delta-deltaCT method for relative quantification and all samples were normalized to a housekeeping gene, Hprt.

### Western blot analysis and antibodies

Cells were lysed with RIPA buffer containing a complete protease and phosphatase inhibitor tablet (Roche #11836153001, #PHOSS-RO). Lysates were cleared by centrifugation at 14,000xq for 15 min at 5°_C. Supernatant fractions were used for Western blot. Protein extracts or purified sEVs were quantified for protein content using the bicinchoninic acid assay (Pierce™ BCA Protein Assay Kit, Thermo Scientific #23225). Equal amounts of cell lysate or purified sEVs were resolved by SDSPAGE and transferred into a polyvinylidene difluoride (PVDF) membrane (Millipore #IPVH00010). After blocking with 5% milk for 1 hour RT, membranes were incubated with the antibody As556 IgG EMILIN-1 (Rabbit Polyclonal from CRO, Italy, 1/2000) overnight at 4°_C. Antibodies to β-actin (Mouse Monoclonal, #A5441 (Sigma), 1/10000) for cells, and Alix 3A9 (Mouse Monoclonal, #2171S (Cell Signaling), 1/1000) for sEVs, were used as loading controls. The intensities of the immunoreactive bands were quantified by densitometry using ImageJ software (NIH).

### Immunofluorescence

Cells were fixed with 4% paraformaldehyde (PFA, Electron Microscopy Sciences) for 20 min at RT, followed by permeabilization with 0.1% Triton X-100 (Sigma #11332481001) in PBS for 10 min RT. After washing with PBS, to avoid antibody unspecific interactions coverslips were incubated with PBS 5% Donkey Serum (Sigma #D9663), 1% BSA, and 0.05% Triton for 45 min at RT and stained with primary antibody As556 IgG EMILIN-1 (Rabbit Polyclonal from CRO, Italy, 1/2000) 4°_C overnight. Then, samples were rinsed and incubated with an Alexa Fluor 488 conjugated secondary antibody (Donkey, Rabbit IgG #A21206 (Life Technologies), 1/200). 40,6-diamidino-2-phenylindole (DAPI) was used for nuclear staining. Digitalized images were generated using a Leica TCS SP5 X AOBS or Leica TCS SP5 AOBS confocal microscopes (63X HCXPLAPO 1.4 N.A) and analyzed using Fiji software.

### GW4869 treatment in vitro

Melanoma cells were treated during 24h with the inhibitor GW4869 (Selleckchem #S7609) at 10 μM per 3×10^5^ of cells seeded in 6-well plates. PBS-DMSO was added as control. After the treatment, cells were stained following the protocol of immunofluorescence previously described. Digitalized images were generated using a Leica TCS SP8 FSU AOBS confocal microscope and analyzed using Fiji software.

### MG-132 treatment *in vitro*

Melanoma cells were treated during 16h with the inhibitor MG-132 (Sigma #1211877-36-9) at 8 μM per 3×10^5^ of cells seeded in 6-well plates. PBS-DMSO was added as control. After the treatment, cells were stained following the protocol of immunofluorescence previously described. Digitalized images were generated using a Leica TCS SP8 FSU AOBS confocal microscope and analyzed using Fiji software.

### Plasmids design and cloning strategies

For generation of HA-EMILIN-1 transfectants, B16-F1 GFP-luc cell line was transfected with pCMV3-N-HA (N-terminal HA-tagged) plasmid (Sinobiological #CV017) in which human EMILIN-1 full sequence cDNA was cloned, control cell line was generated using the empty vector pCMV3-N-HA. The cloning and primers designed for the generation of the vector and the insert fragments sharing overlapping were done following Gibson Assembly NEB protocol (NEB #E2611S/L) and SnapGene Software. For transfection experiments we used the Lipofectamine 2000 Transfection Reagent (Thermo Fisher #11668019). The transfection was done in suspension, where 5×105 B16-F1R2 GFP-luc cells were seeded in a T6 multi-well with 8 μL of Lipofectamine reagent and 8 μg of DNA (ratio 1:1 according to manufacturer’s protocol). 16h later, medium was removed and fresh medium as added. Neomycin (G-418 Sigma #G8168) selection were added 48h later at 2 mg/ml and 500 μg/ml, during 14 days. Stable transfected clones were isolated from the selected cells using “cloning cylinders” (Sigma #CLS31668) and tripsinization. The primers used for cloning were: EMILIN-1 Fw 5’-TGGAGCTCTGGCTTATCCTTACGACGTGCCTGACTACGCCatggccccccgcaccctctg-3’, EMILIN-1 Rv 5’-GAGGGGCAAACAACAGATGGCTGGCAACTAGAAGGCACAGctacgcgtgttcaagctctggg-3’, bGH (poly A) Fw 5’-CTGTGCCTTCTAGTTGCCAGCC-3’, HAtag Rv 5’-GGCGTAGTCAGGCACGTCGTA-3’. Two different polymerases were used due to the length of the vector. Platinium SuperFi DNA polymerase (Invitrogen #12351010) was used for vector amplification following the 3-steps protocol (<10 kb) and Platinum™ Pfx DNA Polymerase (Life Techlogies #11708021) was used for the insert. The amount of fragments used for assembly was 100 ng of the vector and 150 ng of EMILIN-1 insert. For generation of R914W mutant, we performed a site-directed mutagenesis following QuickChange II Sited-Directed Mutagenesis protocol (Aligent Technologies #200523). The primers used for site-directed mutagenesis were: R914W Fw 5’-AAGTGGAGGCCGTGCTGTCCTGGTCCAACCAGGGCGTGGCCCGC-3’, R914W Rv 5’-GCGGGCCACGCCCTGGTTGGACCAGGACAGCACGGCCTCCACTT-3’.Cells were transformed and positive clones were confirmed by DNA sequencing.

### Cell viability assay

Luminescent Cell Viability Assay, CellTiter-Glo, (Promega #G7570) at different time points (24, 48, 72 h) following manufacturer protocol. The CellTiter-Glo® Luminescent Cell Viability Assay is a method to determine the number of viable cells in culture based on quantitation of the ATP present, which signals the presence of metabolically active cells. Cells were seeded into T96-well plate and luminescence was measured at X 490 nm in a VICTOR Multilabel Plate Reader.

### Cell Cycle

Cell cycle histograms for bulk DNA staining (PI), after addition of EdU, from B16-F1-HA and B16-F1-HA-E1 model were performed at 24, 48, 72, 96 h and 1 week following manufacture protocol (Invitrogen #C10337). Percentage of B16-F1-HA and B6-F1-HAE1 cells in S phase was calculated. The modified thymidine analogue EdU was added 30 min before cell fixation. Cells were fixed by adding 100 μl PFA 4% (in PBS, freshly prepared) and Streptavidin-AF647 (Vector) was used after EdU detection mix step. Data were acquired on BD FACS Canto II, at least 5,000 single alive events were acquired and all data was analyzed using FlowJo software v10 (TreeStar).

### Cell tracking and motility assay

Cell tracking and motility analysis were performed overnight in chamber slide with a IBIDI μ-Slide 8 Well (#80826). Videos were acquired in DM6000B Widefield microscope 20X HCXPLAPO 0.7 N.A. (Leica Microsystems). For each cell model it has been required a cell identification and trajectory reconstruction. It has been performed well on live-cell, time-lapse, phase contrast video microscopy of hundreds of cells in parallel. Three or four positions were selected from the videos acquired.. Cell position, distance and velocity were measured and analyzed from each position selected by ImageJ software.

### Xenograft studies

All animal experiments were performed according to protocols approved by the Institutional Ethics Committee for Research and Animal Welfare (CEIyBA) of the CNIO, the Instituto de Salud Carlos III (CNIO-ISCIII) and the Comunidad Autónoma de Madrid (CAM).

### Tumor growth and metastasis studies

8 to 10 week-old *C57BL/6J.OlaHsd* male mice were injected in the flank with 1×10^6^ melanoma cells. Tumor volume was monitored 2-3 times per week. Animals were sacrificed when tumor volume reach 1200 mm3.1200 mm3. To analyze the metastatic spread through the lymphatic system, 8 to 10 week-old C57BL/6J.OlaHsd male mice were injected intra-footpad with 2×104 with melanoma cells. Animals were sacrificed 21 days after injection. Luciferase imaging was done ex vivo using the IVIS Spectrum system in both approaches. Popliteal lymph nodes were paraffin embedded and stained with HMB45, percentage of melanoma positive cells quantification was also performed.

### *In vivo* Imaging System

Luciferase imaging was performed using the IVIS Spectrum system (Caliper, Xenogen). Tumor bearing mice were anesthetized (using isoflurane 3-4% and 0,5% O_2_), and D-luciferin (50 mg kg−1 in 100 μl PBS) was administered. Eight minutes later, mice were euthanized and their organs were analyzed for luciferase expression. Data were quantified with Living Imaging software 4.7.2.

### Histological studies

Tissue samples were fixed in 10% neutral buffered formalin (4% formaldehyde in solution), paraffin-embedded and cut at 3 μm, mounted in superfrost®plus slides and dried overnight. For different staining methods, slides were deparaffinized in xylene and re-hydrated through a series of graded ethanol until water. Consecutive sections were stained with hematoxylin and eosin (H&E), and several immunohistochemistry reactions were performed in an automated immunostaining platform (Autostainer Link 48, Dako; Ventana Discovery XT, Roche). Antigen retrieval was first performed with the appropriate pH buffer, (Low pH buffer, Dako; CC1m, Ventana, Roche) and endogenous peroxidase was blocked (peroxide hydrogen at 3%). Then, slides were incubated with the appropriate primary antibody As556 IgG EMILIN-1 (Rabbit Polyclonal from CRO, Italy). After the primary antibody, slides were incubated with the corresponding secondary antibodies. Immunohistochemical reaction was developed using 3, 30-diaminobenzidine tetrahydrochloride (DAB) or Purple Kit (Chromo Map DAB or Purple Kit, Ventana, Roche; DAB (Dako) and nuclei were counterstained with Carazzi’s hematoxylin. Finally, the slides were dehydrated, cleared and mounted with a permanent mounting medium for microscopic evaluation. Positive control sections known to be primary antibody positive were included for each staining run. Intensity of Emilin-1 expression in tumors was analyzed and scanned using ZEISS ZEN Microscope software, intensity above average was considered as high expression and below average as low.

### Statistical analyzes

Error bars in the graphical data represent means ± s.e.m. Mouse experiments were performed using at least three mice per treatment group. P values of P < 0.05 were considered statistically significant by Student’s t test or ANOVA. For the tumor growth analyzes, we performed two-way ANOVA statistical analyzes using GraphPad Prism software.

---

**Supplementary Fig.1**. Characterization of ES-derived EVs. Size distribution of particles determined by NTA and the concentration of particles derived from B16-F1, B16-F1R2 and B16-F10 mouse melanoma models. Error bars indicate + / −1 standard error of the mean.

**Supplementary Fig.2**. Principal complex networks visualized with ClueGO (Cytoscape plug-in) from the large cluster of genes obtain after proteomic and RNA sequencing data integration. A) KEGG and B) Reactome Pathways Gene Set up-regulated. Enrichment/Depletion (two-side hypergeometric test) Bonferroni step down.

**Supplementary Fig.3**. Analysis of the effect of MG-132. A) Analysis of EMILIN-1 (in green) expression and localization by confocal immunofluorescence (scale 18 μm) in melanocytes, B16-F1 and B16-F1R2 cell lines before and after the treatment with 8 μM MG-132 during 16 hours. Cell nuclei were stained with DAPI (in blue) B) quantification of EMILIN-1 expression (green signal) in C), *p<0.05 by Non-parametric t-test.

