## Supplementary figures and images for "Inactivation of EMILIN-1 by proteolysis and secretion in small extracellular vesicles favors melanoma progression and metastasis"

### Supplementary Figure 1

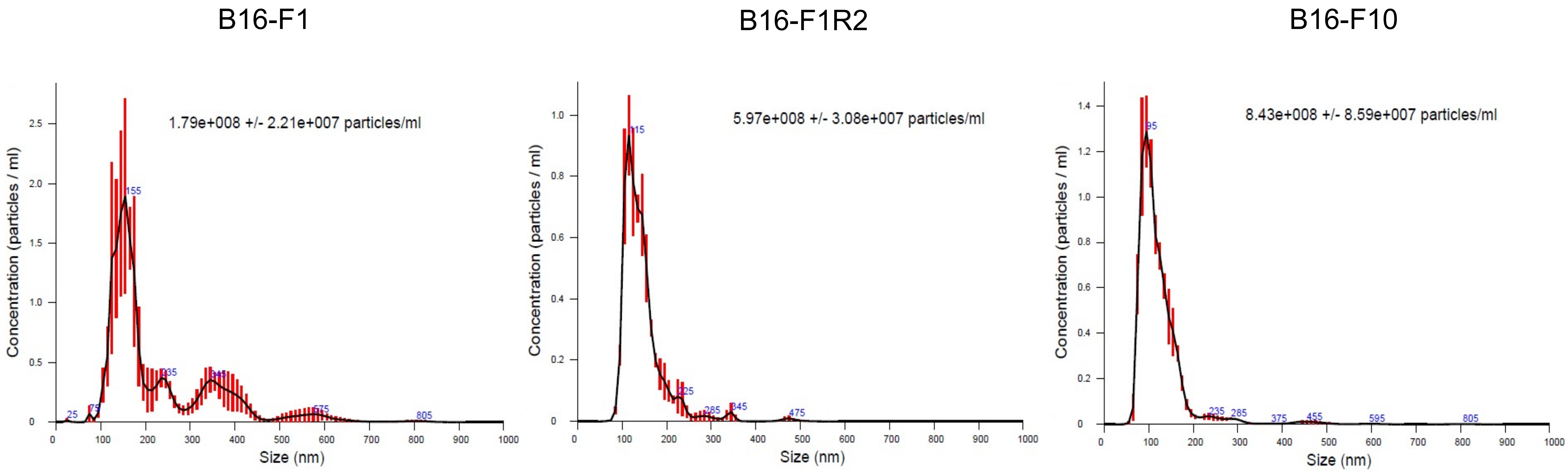


**Supplementary Figure 1**

### Supplementary Figure 2

A)

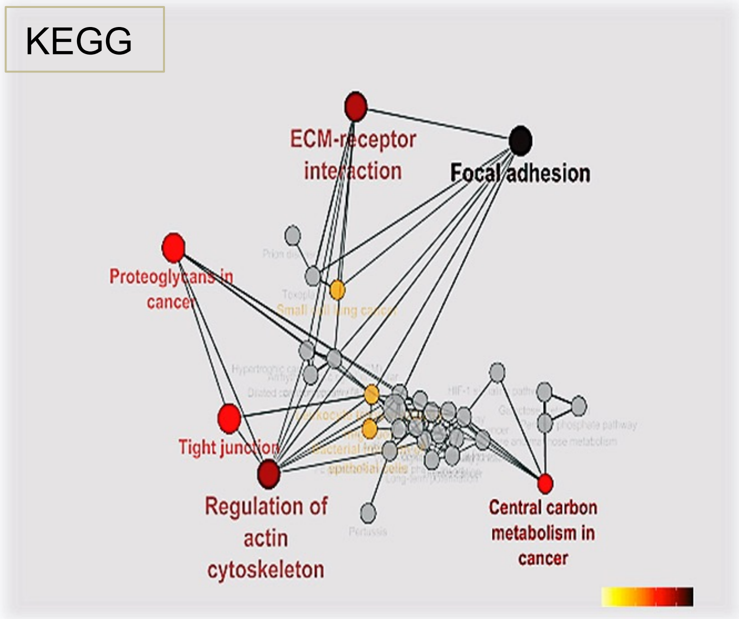


B)

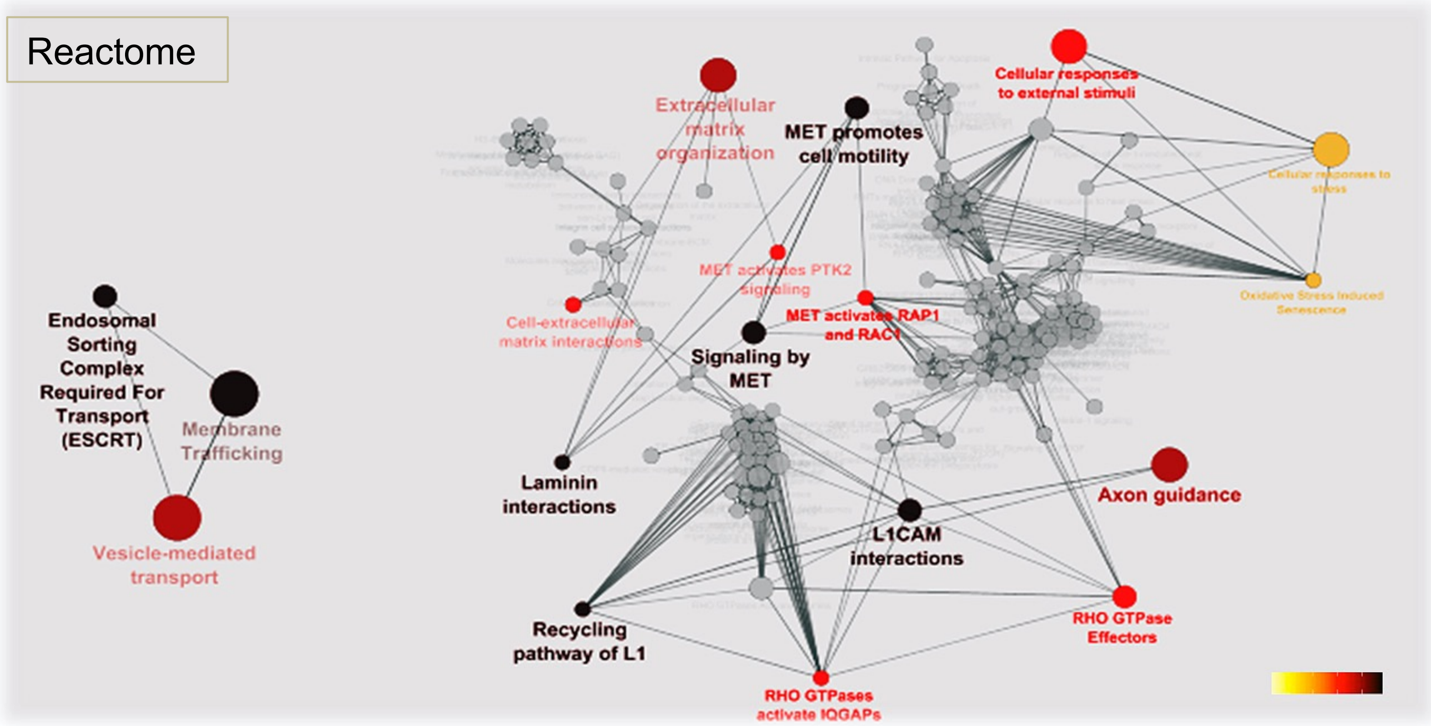


**Supplementary Figure 2**

### Supplementary Figure 3

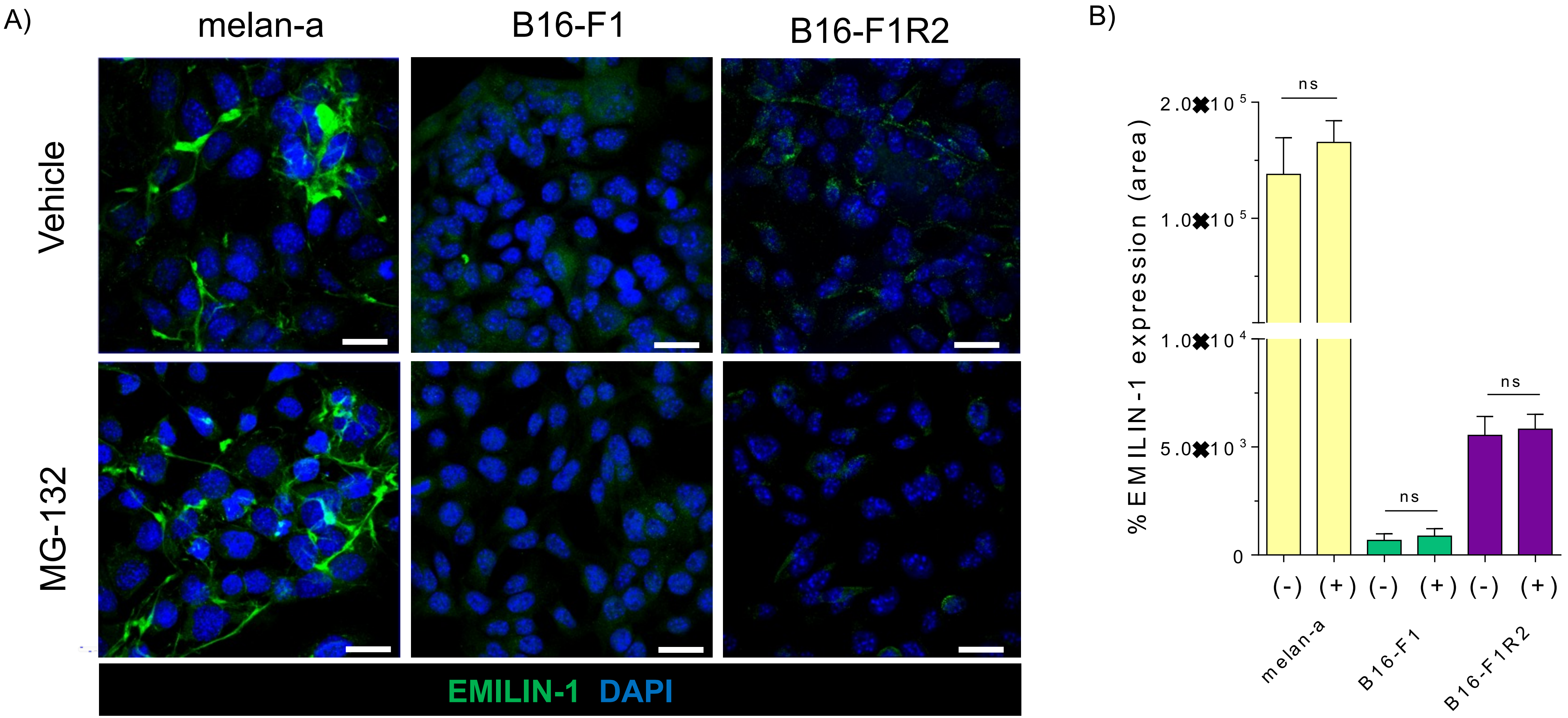


**Supplementary Figure 3**
