## Supplementary Table 1 for "Inactivation of EMILIN-1 by proteolysis and secretion in small extracellular vesicles favors melanoma progression and metastasis"

**Supplementary Table 1.** Top genes most upregulated in B16-F1R2 model compared to B16-F1.

| **Gene Symbol** | **SPOT** | **B16-F1** | **B16-F1R2** |
| --- | --- | --- | --- |
| **LGALS7** | **ID_11499** | **0,12** | **142,72** |
| **CRABP2** | **ID_5108** | **1,95** | **116,7** |
| **RAB7L1** | **ID_17830** | **16,11** | **91,8** |
| **CLU** | **ID_4807** | **0,14** | **72,66** |
| **HSPB1** | **ID_10189** | **15,85** | **67,63** |
| **ACSBG1** | **ID_1937** | **0,26** | **61,1** |
| **CHCHD7** | **ID_4545** | **19,62** | **52,66** |
| **SLC40A1** | **ID_19639** | **1,82** | **52,08** |
| **S100A6** | **ID_18726** | **4,46** | **51,7** |
| **MDK** | **ID_12172** | **0,54** | **46,82** |
| **CSRP2** | **ID_5237** | **17,28** | **46,22** |
| **GRHPR** | **ID_9467** | **15,04** | **45,76** |
| **CCL5** | **ID_4086** | **6,44** | **43,31** |
| **GSTA4** | **ID_9546** | **1,42** | **40,47** |
| **EMILIN1** | **ID_6640** | **4,45** | **40,44** |
| **PVRL2** | **ID_17708** | **19,94** | **39,96** |
| **FAM210B** | **ID_7110** | **8,49** | **38,97** |
| **WFIKKN2** | **ID_23159** | **0,34** | **38,88** |
| **MYC** | **ID_13939** | **17,64** | **38,43** |
| **KDM1B** | **ID_10928** | **4,59** | **38,4** |
| **IL17B** | **ID_10424** | **0,1** | **38,26** |
| **ESPN** | **ID_6830** | **12,97** | **37,94** |
| **PDLIM2** | **ID_16447** | **3,36** | **35** |
| **BNIP3** | **ID_3442** | **11,75** | **34,77** |
| **TMEM56** | **ID_21570** | **11,03** | **33,88** |
| **ISG15** | **ID_10642** | **9,54** | **33,2** |
| **CRMP1** | **ID_5155** | **7,04** | **32,39** |
| **LIMD2** | **ID_11543** | **14,1** | **30,05** |
| **S100A16** | **ID_18721** | **1,12** | **29,94** |
| **SLC35G1** | **ID_19600** | **12,38** | **29,34** |
| **FN1** | **ID_7515** | **4,85** | **29,26** |
| **ZDHHC2** | **ID_23426** | **13,09** | **27,76** |
| **SPHK1** | **ID_20260** | **7,23** | **27,55** |
| **APOBEC2** | **ID_2585** | **0,46** | **27,55** |
| **DAB2** | **ID_5635** | **2,24** | **27,15** |
| **ADSSL1** | **ID_2128** | **3,44** | **27,13** |
| **MALAT1** | **ID_11939** | **12,77** | **27,12** |
| **ATP1A3** | **ID_2953** | **0,11** | **26,68** |
| **MCOLN3** | **ID_12149** | **6,73** | **26,47** |
| **RCN3** | **ID_18067** | **0,64** | **26,21** |
| **KDELR3** | **ID_10925** | **9,98** | **26,16** |
| **CD72** | **ID_4230** | **3,12** | **25,75** |
| **GSTK1** | **ID_9548** | **0,69** | **25,48** |
| **GPT** | **ID_9435** | **5,13** | **25,17** |
| **PTGIR** | **ID_17615** | **0,3** | **24,55** |
| **RGS10** | **ID_18182** | **11,77** | **24,1** |
| **LRP5** | **ID_11681** | **7,34** | **23,84** |
| **FKBP10** | **ID_7464** | **1,49** | **23,54** |
| **SLC27A6** | **ID_19538** | **6,14** | **23,53** |
| **MMP2** | **ID_13522** | **1,14** | **22,81** |
| **AK1** | **ID_2219** | **4,54** | **22,29** |
| **CORO2B** | **ID_5018** | **9,21** | **22,06** |
| **RAB17** | **ID_17780** | **4,24** | **22** |
| **TMOD1** | **ID_21616** | **8,07** | **21,97** |
| **EGFL7** | **ID_6501** | **10,48** | **21,43** |
| **HMGN5** | **ID_10003** | **10,46** | **20,92** |
| **FAM3B** | **ID_7143** | **0,06** | **20,84** |
| **NXN** | **ID_14757** | **7,37** | **20,61** |
| **STEAP1** | **ID_20525** | **4,71** | **20,13** |
