## Supplementary Table 2 for "Inactivation of EMILIN-1 by proteolysis and secretion in small extracellular vesicles favors melanoma progression and metastasis"

**Supplementary Table 2.** Integration of the mass spectrometry data (proteins secreted in sEVs) with the RNA sequencing data.

| **Gene names** | **PROT log2 (R2 / F1)** | **RNA SEQ log2 (R2 / F1)** |
| --- | --- | --- |
| **Clu** | **6,76** | **6,02** |
| **Acsbg1** | **6,20** | **5,62** |
| **Mdk** | **3,88** | **4,95** |
| **Gdf15** | **4,14** | **4,39** |
| **Hapln1** | **3,30** | **4,36** |
| **Slc40a1** | **1,73** | **4,23** |
| **Cd37** | **4,65** | **4,17** |
| **Slc12a8** | **1,34** | **4,12** |
| **Icam2** | **3,20** | **3,27** |
| **Casq1** | **6,81** | **3,22** |
| ***Emilin1*** | **4,00** | **2,92** |
| **Igf2bp3** | **1,32** | **2,86** |
| **Mbp** | **2,79** | **2,54** |
| **Slc38a3** | **1,77** | **2,49** |
| **Col5a1** | **3,61** | **2,39** |
| **Pdgfrb** | **2,91** | **2,24** |
| **Glipr2** | **1,77** | **2,19** |
| **Rab17** | **1,51** | **2,13** |
| **Crmp1** | **1,99** | **2,05** |
